# Aquaporin mediating stomatal closure is associated with water conservation under mild water deficit

**DOI:** 10.1101/2020.04.15.042234

**Authors:** Lei Ding, François Chaumont

## Abstract

- Contradictory results indicate that aquaporins might facilitate the diffusion of both water and H_2_O_2_ during abscisic acid (ABA) triggered stomatal closure. Here, we tested whether maize plasma membrane PIP2;5 aquaporin regulates stomatal closure under water deficit or ABA treatment in intact plants, detached leaves, and peeled epidermis.
- Transpiration, stomatal conductance and aperture, as well as reactive oxygen species (ROS) in stomatal complexes were studied in maize lines deregulated in *PIP2;5* gene expression, under water deficit and/or ABA treatments.
- In well-watered conditions, the PIP2;5 overexpressing (OE) plants transpired more than the wild-type plants (WT), while no significant difference in transpiration was observed between *pip2;5* KO and WT plants. Upon mild-water deficit or low ABA concentration treatment, the transpiration and stomatal conductance decreased more in PIP2;5 OE, and less in *pip2;5* KO lines, in comparison with WT plants. Using isolated epidermis, ABA treatment induced faster stomatal closing in PIP2;5 OE lines compared to the WT, while *pip2;5* KO stomata were ABA insensitive. These phenotypes were associated with guard cell ROS accumulation.
- Together, these data indicate that maize PIP2;5 regulates early stomatal closure for water conservation upon a water deficit environment.

## Introduction

Stomata are micropores in the leaf epidermis formed by two guard cells allowing CO_2_ uptake for photosynthesis at the expense of water loss by transpiration. The kinetics and magnitude of stomatal aperture/conductance are regulated by natural environmental fluctuations, such as light, soil water availability, CO_2_, and air vapor pressure deficit (VPD) (Tardieu & Simonneau, 1998; Zhang *et al.*, 2018; Sussmilch *et al.*, 2019). With a rapid stomatal movement response to the changing environment, carbon assimilation, water use efficiency, and plant growth can be improved (Papanatsiou *et al.*, 2019). In dry environments, including dry soil and atmosphere, stomatal closure is mediated by abscisic acid (ABA) signaling in guard cells (Cai *et al.*, 2017; Pantin & Blatt, 2018; Sussmilch *et al.*, 2019) and bundle sheath cells (Pantin *et al.*, 2013; Moshelion *et al.*, 2015). In the latter cells, ABA signaling triggers a decrease in leaf hydraulic conductance (K_leaf_) and leaf water potential (Ψ_leaf_), contributing to stomatal closure via a hydraulic mechanism (Pantin *et al.*, 2013). In stomata, ABA signaling activates the membrane transport system, including potassium and anion channels, and possibly aquaporins, to decrease guard cell turgor pressure and volume, leading to stomatal closure (Grondin *et al.*, 2015; Jezek & Blatt, 2017; Papanatsiou *et al.*, 2019).

For many years, the roles of aquaporins in controlling stomatal movement remained hypothetical, even if their role in facilitating membrane diffusion of water and small solutes makes them putative important actors for guard cell turgor pressure adjustment (Chen *et al.*, 2017; Hachez *et al.*, 2017; Jezek & Blatt, 2017; Ding & Chaumont, 2020). In 2015, the first direct evidence highlighted the involvement of *Arabidopsis* PIP2;1 in ABA induced stomatal closure, the closure being impaired in *pip2;1* KO lines (Grondin *et al.*, 2015). In addition, ABA-activated OST1 kinase phosphorylates the Ser-121 of PIP2;1, an event known to activate PIP aquaporin water channel activity (Grondin *et al.*, 2015). PIP2;1 also facilitates H_2_O_2_ diffusion into guard cells to mediate ABA- and pathogen-triggered stomatal closure (Rodrigues *et al.*, 2017). At the same time, PIP2;1 was suggested to be involved in stomatal closure induced by greater CO_2_ concentrations (Wang *et al.*, 2016). However, the ABA-induced stomatal closure phenotype of the *Arabidopsis pip2;1* KO line could not be reproduced, and even a quadruple *pip1;1*, *pip1;2*, *pip2;1*, and *pip2;2* mutant did not exhibit any difference in stomatal closure and conductance (Wang *et al.*, 2016; Ceciliato *et al.*, 2019). Further evidence is required to ascertain the exact role(s) of aquaporins in stomatal movement in other species (Ding & Chaumont, 2020).

In this study, we examined whether *PIP2;5* gene expression deregulation affects maize stomatal closure induced by water deficit and ABA treatments in intact plants, detached leaves, and peeled epidermis. We previously showed that PIP2;5 is a main actor in controlling cell and tissue hydraulic conductivity, as well as in regulating plant growth under water deficit conditions (Hachez *et al.*, 2012; Ding *et al.*, 2020). Additionally, H_2_O_2_ membrane diffusion is enhanced by expressing PIP2;5 in yeast (Bienert *et al.*, 2014). Those results indicate maize PIP2;5 might be involved in stomatal movement regulation. Here, we report that PIP2;5 regulates early stomatal closure and transpiration decrease upon water deficit. This data is of potential interest to design new strategies for crop breeding and enhance abiotic stress resilience and productivity in changing climate scenarios.

## Materials and Methods

### Plant materials and growth conditions

Two PIP2;5 OE and one *pip2;5* KO maize lines used in this study were described in. The heterozygous plants were self-pollinated and, in the next generation, the homozygous plants were identified by quantitative real-time PCR (qRT-PCR) (PIP2;5 OE lines), and by PCR on genomic DNA (*pip2;5* KO lines). Homozygous plants segregating for the *PIP2;5* transgene (PIP2;5 OE) and non-transgenic siblings (WT-B104) or *pip2;5* KO and siblings without the Mu transposon (WT-W22) were utilized. Plants were grown in soil in a growth chamber, with a 16 h/8 h light/dark cycle (25/18°C) and a daytime light intensity of 200 μmol.m^−2^.s^−1^ at the leaf top. The chamber humidity during the light cycle was 50%∼70%.

### Whole plant transpiration measurements

Whole plant transpiration was determined by gravimetry. Briefly, three-week-old maize plants were placed on a balance (FX-3000i, A&D, Japan) and the weight was recorded every 10 min with a weighing data logger (AD-1688, A&D, Japan) in the light/dark cycle. After measurement, the leaf was scanned and the leaf area was analyzed with ImageJ software. Whole plant transpiration was calculated with the weight loss and normalized with the leaf area. Soil water potential was measured by WP4C DewPoint Potentiometer (Decagon Devices, Inc, Pullman, WA, US).

### Measurements of g_s_

g_s_ was measured with a Li-Cor 6400XT portable photosynthesis system (Li-Cor Inc., Lincoln, NE, US). The third leaf from two-week-old maize plants was excised with ∼3 cm leaf sheath, which was recut inside Milli-Q water to have a final leaf with ∼2 cm leaf sheath. Then, the excised leaf was transferred to a 5-mL Eppendorf tube filled with milli-Q water. This transfer was performed in water to avoid any air entering into the leaf. The leaf was stabilized under light for ∼30 min and then placed in the Li-Cor 6400XT system leaf chamber. During the measurement, the chamber temperature was maintained at ∼28°C and the light intensity was 1500 μmol m^−2^ s^−1^ (with 10% blue + 90% red light). CO_2_ concentration was set at 400 μmol mol^−1^ controlled with a CO_2_ cartridge (Liss, Hungary). The relative humidity was 40%∼50%. g_s_ was recorded every 30 s for ∼20 min before ABA was added, and then for another ∼30 min with the indicated ABA concentrations.

### Stomatal aperture measurements

Stomatal aperture was measured from 4^th^ leaf abaxial epidermal peels from three-week-old maize plants. Firstly, the fully expanded leaf was excised and cut into ∼2 cm long sections, with the abaxial side floated on the incubation buffer (10 mM KCl, 50 μM CaCl_2_, 10 mM MES, pH 5.6 (Tris)) (Gao *et al.*, 2017) (Greiner Bio-One, Vilvoorde, Belgium). The leaves were exposed to light (∼200 μmol m^−2^ s^−1^) for 2 h to induce stomatal opening. Then, the leaf epidermis was peeled and the strips were floated on the incubation buffer containing either 10 μM ABA (Sigma-Aldrich, US) (30 mM ABA stock in ethanol) or ethanol as a control. Stomatal aperture was checked after 30, 60, and 120 min treatments with a fluorescence microscope (Axio observer 7, Zeiss, Germany) equipped with a CCD camera (ORCA-Flash 4.0 LT C11440, Hamamatsu, Japan). The aperture size was measured with ImageJ software.

### ROS measurements in stomata

ROS in stomata was detected by H_2_DCFDA staining (Pei *et al.*, 2000; Iwai *et al.*, 2019). Stomatal opening was induced as above. After 2 h of induction, the epidermis was peeled inside the incubation buffer, and the strip was submerged into the incubation buffer with 10 μM ABA. After the indicated time, the epidermal strip was washed three times with Milli-Q water and stained with 50 μM H_2_DCFDA (Sigma-Aldrich) (10 mM stock in DMSO) in incubation buffer for 20 min in the dark. The strip was washed three times and the fluorescence signal was detected by microscopy (excitation: 494 nm; emission: 517 nm) and quantified with ImageJ software.

### Statistical analyses

Statistical analyses were performed with GraphPad Prism 5 (GraphPad Software). Student’s *t*-test was applied to determine the significant differences in Fig. 1 and Fig. 2. One-way ANOVA with Tukey post-test was applied to compare the significant differences in Fig. 3 and Fig. 4.

**Fig.1.**
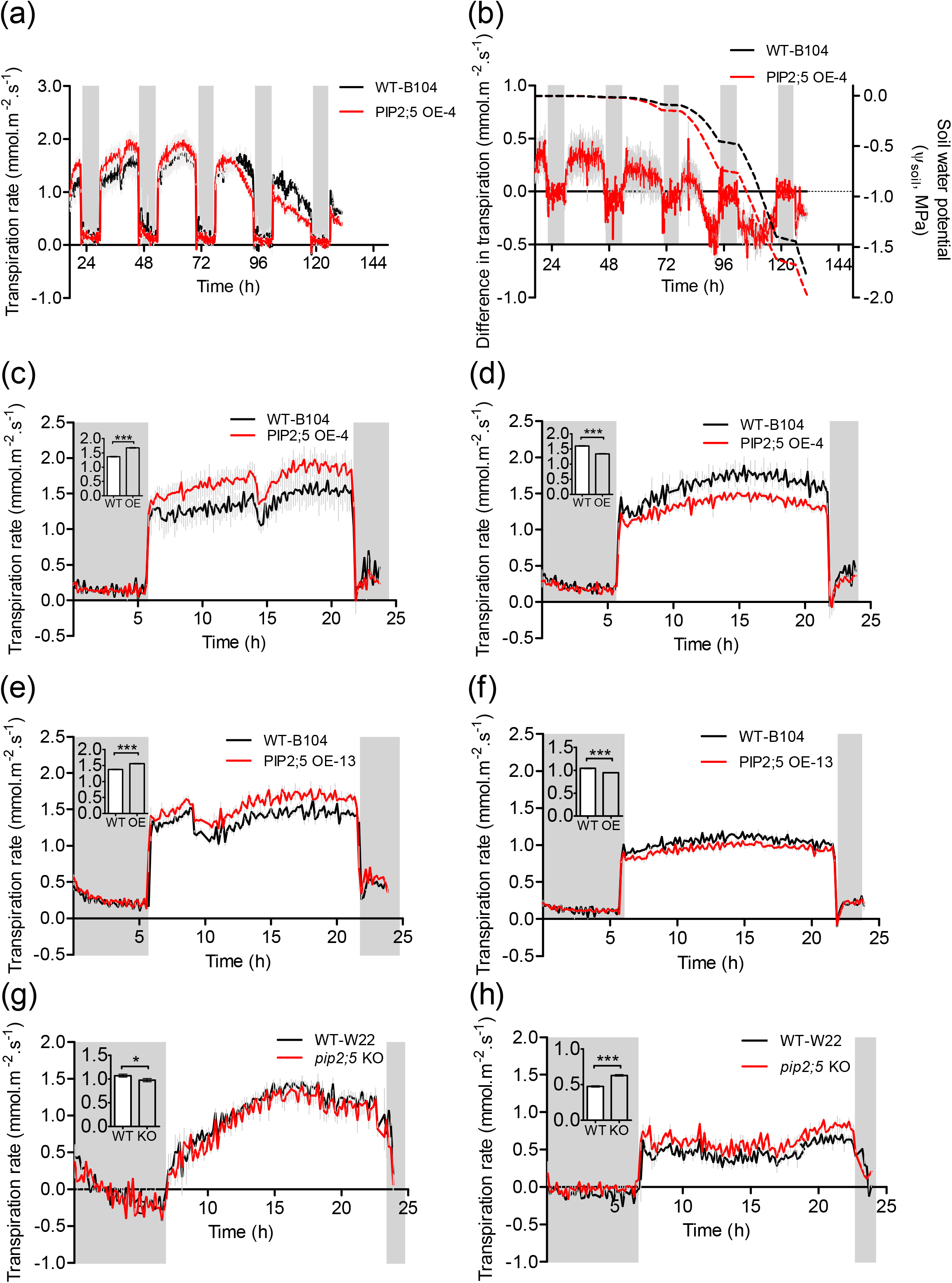
Transpiration in maize lines deregulated in *PIP2;5* gene expression. (a) Transpiration of WT-B104 and PIP2;5 OE-4 plants were recorded continuously from well watering to water deficit by stopping plant watering. (b) The transpiration difference between PIP2;5 OE-4 and WT-B104 plants. The soil water potential was indicated by the dashed lines. (c-h) Transpiration was recorded continuously for 24 h under well-watered conditions (c, e, g) and mild soil water deficit (d, f, h). (c) and (d) PIP2;5 OE-4 lines. (e) and (f) PIP2;5 OE-13 lines. (g) and (h) *pip2;5* KO lines. The insets indicate the day mean transpiration rate. The PIP2;5 OE, *pip2;5* KO, and corresponding WT plants are indicated by OE, KO, and WT, respectively. The dark period is indicated by the grey bar. The soil water potential was ∼−0.10 MPa in mild soil water deficit treatment. The mean values were from two to four plants. The error bars indicate the standard error (SE). Student’s *t*-test was applied to compare the significant difference of day mean transpiration between the lines at levels of p<0.001 (***) and p<0.05 (*).

**Fig.2.**
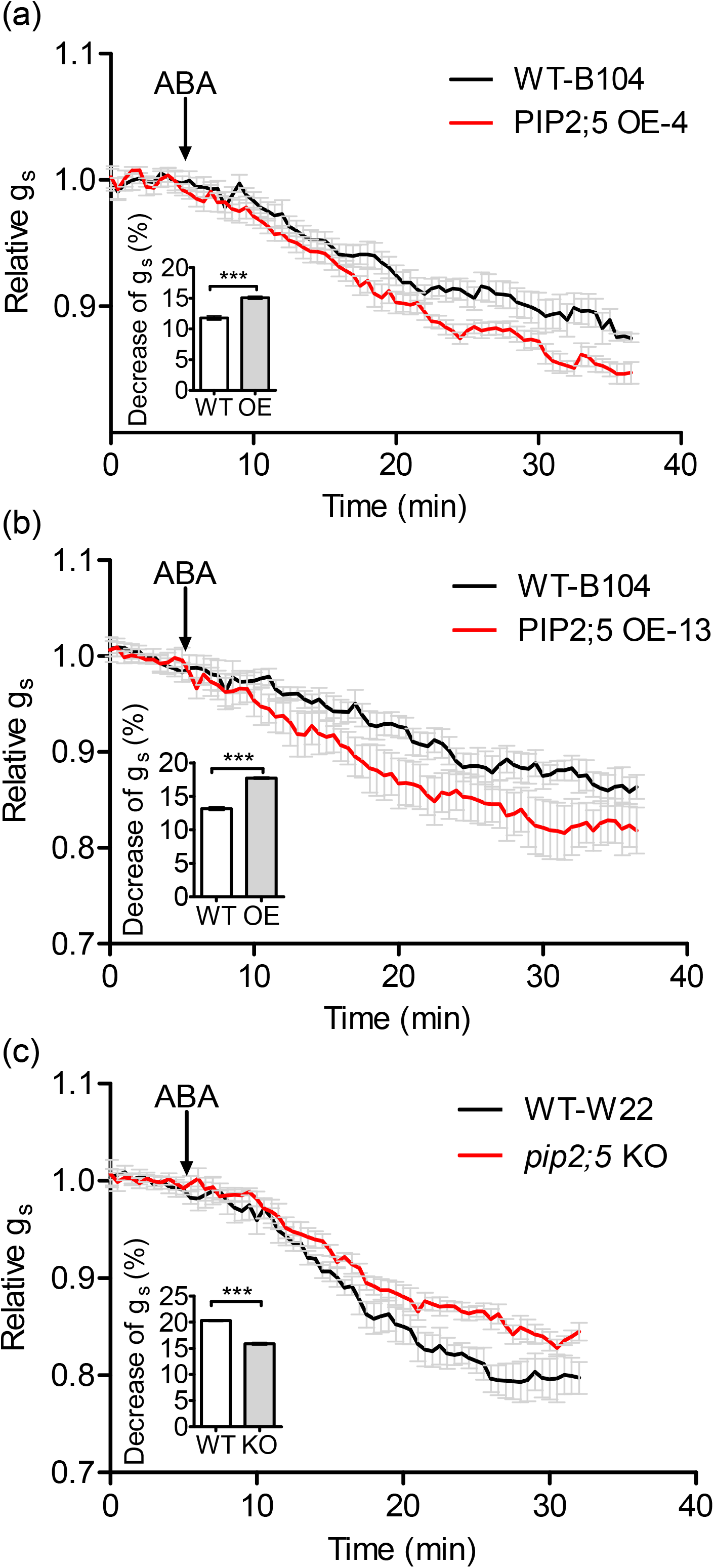
g_s_ dynamics after ABA treatment. (a) PIP2;5 OE-4. (b) PIP2;5 OE-13. (c) *pip2;5* KO. The black arrows indicate the time of 0.1 μM ABA addition. The mean g_s_ values were calculated during 5 min before ABA addition, and the data were normalized with these mean values. The data was calculated from 5-9 leaves in two independent experiments. The insets indicate the g_s_ percentage decrease, calculated from the measurement last 5 min. The student’s *t*-test was applied to compare the significant difference in g_s_ decrease between the lines at the p<0.001(***) level.

**Fig.3.**
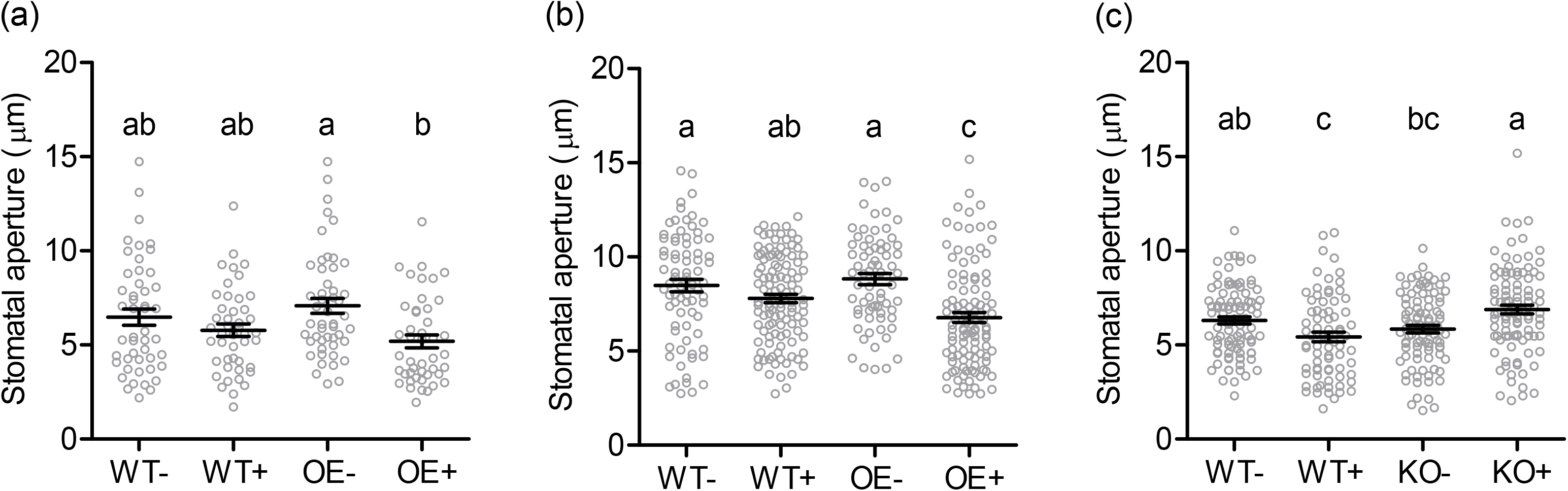
Stomatal aperture after ABA treatments. (a) PIP2;5 OE-4. (b) PIP2;5 OE-13. (c) *pip2;5* KO. The epidermal peels were incubated in 10 μM ABA for 30 min, and stomatal aperture was measured from n > 40 stomata in two independent experiments. Data are expressed as the mean ± SE and individual data points are shown (opened circles). The PIP2;5 OE, *pip2;5* KO, and the corresponding WT plants are indicated by OE, KO, and WT, respectively. “−” and “+” indicate with and without ABA treatments, respectively. One-way ANOVA with Tukey post-test was applied to compare the stomatal aperture significant difference between ABA and the control treatments at the p<0.05 level. The significant difference is indicated by different letters.

**Fig.4.**
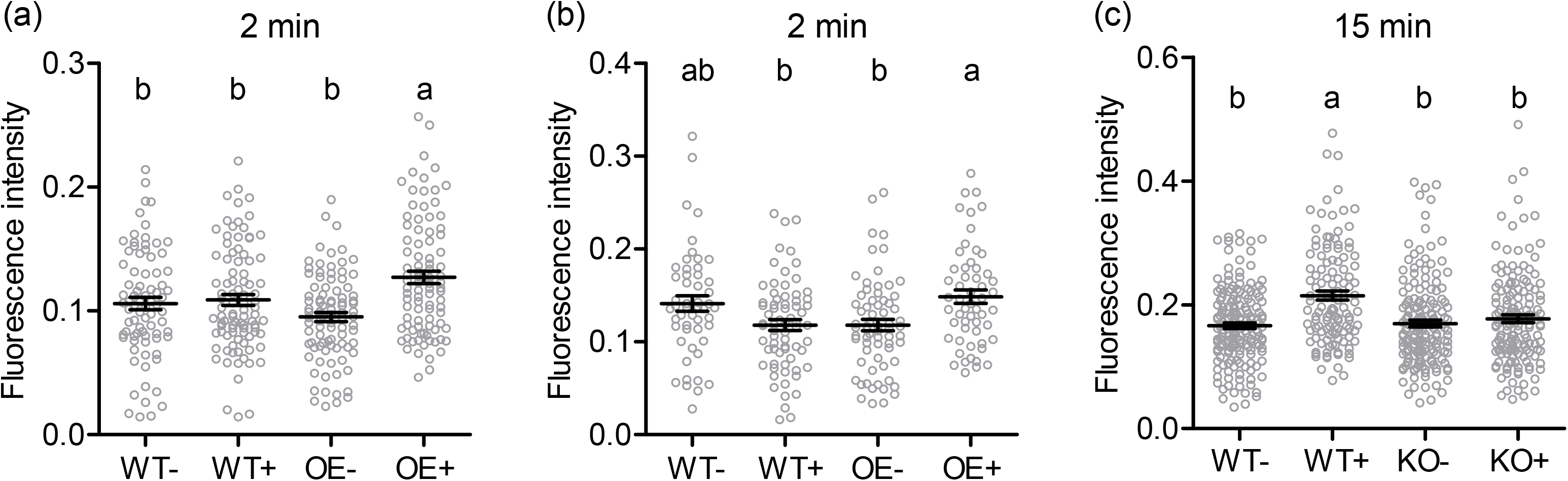
ROS accumulation in guard cells after ABA treatments. (a) PIP2;5 OE-4. (b) PIP2;5 OE-13. (c) *pip2;5* KO. The epidermal peels were incubated in 10 μM ABA for 2 min in (a) and (b), for 15 min in (c), and the H_2_DCFDA signal was detected. The data was calculated from n>50 guard cells in two to three independent experiments. The PIP2;5 OE, *pip2;5* KO, and the corresponding WT plants are indicated by OE, KO and WT, respectively. “−” and “+” indicate with and without ABA treatments, respectively. One-way ANOVA with Tukey post-test was applied to compare the stomatal aperture significant difference between ABA and the control treatments at the p<0.05 level. The significant difference is indicated by different letters.

## Results

### *PIP2;5* gene expression deregulation affects transpiration

Whole plant transpiration was monitored continuously for two PIP2;5 OE (PIP2;5 OE-4 and PIP2;5 OE-13) and one *pip2;5* KO lines, together with the respective WT siblings (WT-B104 and WT-W22) (Ding *et al.*, 2020), under well-watered conditions and soil water deficit. We first measured transpiration with PIP2;5 OE-4 and WT siblings (WT-B104) grown in progressively drying soil conditions (Fig. 1a, b). In well-watered conditions, a greater transpiration was recorded in PIP2;5 OE-4 plants compared with the WT plants. However, the transpiration rate of PIP2;5 OE plants was more sensitive to very mild soil water deficit conditions (Ψ_soil_ ≈ −0.05 MPa), indicated by the decrease in the difference in transpiration between the OE and WT-B104 plants (Fig. 1b).

We compared the transpiration in well-watered (Ψ_soil_ ≈ 0 MPa) and mild water deficit conditions (Ψ_soil_ ≈ −0.10 / −0.20 MPa) for all the lines. In well-watered conditions, the transpiration of PIP2;5 OE-4 and OE-13 plants was 22% and 14% greater during the day, respectively, compared with WT-B104 (Fig. 1c, e). No significant transpiration difference was observed between *pip2;5* KO and WT-W22 plants (Fig. 1g). On the other hand, under mild water deficit, PIP2;5 OE-4 and OE-13 plants transpired 16% and 9% less water during the day, respectively, in comparison with WT-B104 (Fig.1d, f; Fig. S1a). In this water deficit condition, the transpiration of *pip2;5* KO plants was 31% greater during the day than the WT-W22 siblings (Fig. 1h). Interestingly, under severe soil water deficit conditions (Ψ_soil_ ≈ −0.3 to −0.5 MPa), PIP2;5 OE plants transpired more than the WT-B104 plants (Fig. S1).

### ABA-induced stomatal closure is altered by *PIP2;5* gene expression deregulation

As transpiration is regulated by g_s_, we investigated whether *PIP2;5* gene expression deregulation affects stomatal behavior after different ABA treatments. We measured g_s_ using excised leaves, and stomatal aperture in epidermal peels under different ABA treatments.

We observed that g_s_ in WT-B104 leaves decreased faster with increasing ABA concentrations (0.1, 0.5, and 10 μM ABA; Fig. S2a). Then we compared g_s_ of WT, PIP2;5 OE, and *pip2;5* KO leaves using 0.1 μM ABA. Interestingly, g_s_ of PIP2;5 OE and *pip2;5* KO leaves decreased faster and slower, respectively, compared with their respective WT siblings (Fig. 2). This difference in g_s_ behavior was not observed when using 0.5 μM ABA (Fig. S2b,c,d).

A significant decrease in stomatal aperture was observed in the peeled epidermis of both PIP2;5 OE lines after 30 min incubation with 10 μM ABA (Fig. 3a,b), while a significant decrease was only observed after 60 min ABA treatment for the WT-B104 or WT-W22 lines (Fig. S3a-d), indicating faster stomatal closure in PIP2;5 OE plants. In contrast, no decrease in stomatal aperture in *pip2;5* KO epidermis upon ABA treatment was recorded whatever the incubation time. Actually, a larger *pip2;5* KO stomatal aperture was recorded after ABA treatment than without treatment (Fig. 3c; Fig. S3e,f), suggesting PIP2;5 has a key role in stomatal ABA sensing.

### ROS accumulation induced by ABA is altered by PIP2;5 deregulation

ABA-induced stomatal closure is dependent on ROS (mainly H_2_O_2_) accumulation in guard cells (Pei *et al.*, 2000; Rodrigues *et al.*, 2017). We followed H_2_O_2_ accumulation in guard cells after epidermis incubation with ABA using the H_2_DCFDA probe. Significant H_2_O_2_ accumulation was measured in PIP2;5 OE guard cells after 2 min ABA (10 μM) incubation, while no accumulation was detected in WT-B104 after this short time (Fig. 4a,b). A significant increase in the H_2_O_2_ signal in WT guard cells was only found after 15 min ABA treatment (Fig. 4c; Fig. S4b,d). On the other hand, no ROS accumulation was observed in *pip2;5* KO guard cells at any ABA incubation time, while the signal increased after 15 min ABA treatment in the WT-W22 siblings (Fig. 4c; Fig. S4e,f).

## Discussion

### PIP2;5 affects transpiration in a water condition dependent manner

Transpiration and g_s_ are regulated by aquaporins expressed in roots and shoots, and transpiration and/or g_s_ changes up to 40% are obtained by aquaporin gene expression manipulation at the whole plant level under controlled growth conditions (Chaumont & Tyerman, 2014; Maurel *et al.*, 2016). In accordance with these observations, the transpiration rate of the PIP2;5 OE lines increased by 14-22% in comparison with the transpiration in WT plants in well-watered soil. This positive effect was possibly due to an increase in tissue hydraulic properties mediated by aquaporins in roots and/or shoots (Shatil-Cohen *et al.*, 2011; Pantin *et al.*, 2013; Prado *et al.*, 2013; Ding *et al.*, 2020). Sade *et al*. (2014a) also found that g_s_ increased in *Arabidopsis* lines overexpressing NtAQP1 (a PIP1 isoform) under the *35S* promoter control or the main photosynthetic tissue promoter *FBPase*, but not when expression was controlled by the stomatal-specific *KST1* promoter. Conversely, transpiration and g_s_ were decreased by constitutively silencing PIP1s, but not by specific silencing in bundle sheath cells (Sade *et al.*, 2014b). Altogether, this data indicates that aquaporins can indirectly affect g_s_ through changes in tissue hydraulic properties affecting signaling processes.

In soil water deficit conditions, the decrease in transpiration and g_s_ results from ABA-induced stomatal closure (Zhu, 2002; Cai *et al.*, 2017). Here, we showed that, in the PIP2;5 OE lines, the transpiration was less in a mild water deficit condition (−0.1 / −0.2 MPa) and g_s_ decreased more rapidly after low ABA concentration (0.1 μM) treatment, compared to WT plants. Interestingly, an inverse behavior was observed for the *pip2;5* KO plants. These results indicate that PIP2;5 is involved in regulating stomatal closure under mild drought stress or ABA treatments. Overexpressing *Vicia faba Vf*PIP1 in Arabidopsis (Cui *et al.*, 2008) or the apple MdPIP1;3 in tomato plants (Wang *et al.*, 2017), also results in faster stomatal closure in transgenic plants than in control plants upon drought stress or ABA treatments. However, this phenotype we reported here appeared to be dependent on the water stress conditions. We showed here that this difference in stomatal closure behavior was only observed under mild soil water deficit or low concentration ABA treatment, but not under severe water deficit or high ABA treatments (Fig. S1; Fig. S2b-d). Those results indicate that stomatal closure is regulated by several factors under drought stress or ABA treatment, involving signaling events both in stomatal complexes and in vascular bundle sheath (Shatil-Cohen *et al.*, 2011; Pantin *et al.*, 2013). In severe soil water deficit conditions (Ψ_soil_ ≈ −0.30∼−0.50 MPa), the PIP2;5 OE maize lines may maintain root water uptake ability and leaf hydraulic conductance (Ding *et al.*, 2020), resulting in similar or even higher plant transpiration and growth as in WT plants. Altogether, these data indicate that the beneficial effect of PIP2;5 overexpression is really dependent on water stress conditions and that, in mild water deficit, PIP2;5 OE lines perform better than control plants by faster stomatal closing and/or increased root water uptake ability, as previously observed for plants overexpressing other PIP aquaporins (Groszmann *et al.*, 2017; Sade & Moshelion, 2017),

### ABA-mediated stomatal closure depends on stomatal expressed PIP2;5

Stomatal aperture and closure depend on signaling molecules such as ABA, H_2_O_2_, and CO_2_, and molecular events leading to variations in guard cell turgor pressure and, thus, on water fluxes across their plasma membrane. Due to their multiple channel specificity, aquaporins are thought to contribute to different stomatal movement processes (Heinen *et al.*, 2014; Chen *et al.*, 2017; Nunes *et al.*, 2020). The first functional evidence of an aquaporin in stomatal movement was provided by studies using epidermal peels from Arabidopsis *pip2;1* KO plants. PIP2;1 was required for ABA-dependent stomatal closure and, during this process, phosphorylation of PIP2;1 by the specific kinase OST1 activated the water and H_2_O_2_ transport (Grondin *et al.*, 2015; Rodrigues *et al.*, 2017). We found that PIP2;5 OE stomata from epidermal peels closed faster upon ABA treatment, while stomatal aperture was insensitive to ABA treatment in *pip2;5* KO plants. This faster stomatal closure was correlated with ROS accumulation (H_2_O_2_) in PIP2;5 OE guard cells. Actually, H_2_O_2_ is a key factor in ABA and CO_2_ signaling regulating stomatal closure (Chater *et al.*, 2015; Rodrigues *et al.*, 2017), and H_2_O_2_, produced in the apoplasm by the activated NADPH oxidases, acts in guard cells as a Ca^2+^ channel activity regulator leading to SLAC1 activation in the plasma membrane (Pei *et al.*, 2000) and stomatal closure. Several aquaporins, including maize PIP2;5, were previously shown to facilitate H_2_O_2_ diffusion when expressed in yeast (Bienert *et al.*, 2007; Bienert *et al.*, 2014) or *in planta* (Tian *et al.*, 2016; Rodrigues *et al.*, 2017). Altogether, these data strongly suggest that PIP2;5 expressed in maize guard cells acts as H_2_O_2_ channels.

Recently, an *Arabidopsis* quadruple *pip1;1*, *pip1;2*, *pip2;1*, and *pip2;2* mutant was generated but did not exhibit any significant difference in stomatal aperture or g_s_ upon 2 μM ABA treatment compared with the WT (Ceciliato *et al.*, 2019). The same group was also unable to confirm the stomatal ABA insensitivity of *pip2;1* mutant plants (Wang *et al.*, 2016). However, these conflicting results might be explained by different plant growth conditions and/or experimental methods. For example, in our study, the decrease or increase in g_s_ observed in PIP2;5 OE plants and *pip2;5* KO plants, respectively, were only recorded using 0.1 μM ABA treatment, but not with 0.5 μM or greater ABA concentrations. This indicated that stomatal closure is due to ABA signaling in guard cells when the epidermis is investigated, while stomatal closure is regulated by ABA signaling in both guard cells and vascular bundle sheath cells in intact leaves (Pantin *et al.*, 2013). We showed that guard cells are more sensitive to mild soil water deficit or low ABA treatment in PIP2;5 OE plants. However, under severe water deficit or high ABA treatment, stomatal closure might be more regulated by ABA signaling in vascular bundle sheath cells, as demonstrated by the fact that high ABA (50 μM) treatment was able to decrease stomata of ABA insensitive mutants (*ost2-1*, *abi1-1*, *slac1-1*), through leaf hydraulic conductance regulation.

In conclusion, our work demonstrates that PIP2;5 is an important player of grass stomatal movement dynamics. It improves ABA signaling sensitivity probably by facilitating H_2_O_2_ accumulation and speeding up stomatal closure. This process is beneficial to plant growth in mild water deficit conditions, allowing water conservation and a greater leaf elongation rate (Ding *et al.*, 2020).

## Acknowledgements

The authors thank Prof. F Van Breusegem for lending the Li-cor6400XT and Robin Pottie for his help with the Li-cor6400XT. We also thank Prof. Mathieu Javaux and Prof. Xavier Draye for lending their water potential meter and balances. This work was supported by the Interuniversity Attraction Poles Programme-Belgian Science Policy (grant IAP7/29), the “Communauté française de Belgique-Actions de Recherches Concertées” (grant ARC16/21-075), and the Pierre and Colette Bauchau Award. L.D. was supported by Incoming Post-doc Move-in Louvain Fellowships co-funded by the Marie Curie Actions, and FSR/UCLouvain researcher funding.

## Author Contribution

L.D and F.C. designed the experiments; L.D. performed the experiments; L.D. and F.C. analyzed the data and wrote the manuscript.

## Supporting information

Fig.S1 Transpiration in PIP2;5 OE plants. (a) The transpiration of PIP2;5 OE-4 plants under mild water deficit and severe water deficit conditions. (b) The transpiration of PIP2;5 OE-13 plants under severe water deficit conditions. The dash lines indicate the soil water potential.

Fig.S2 g_s_ dynamic after ABA treatment. (a) g_s_ responding to different ABA concentrations in WT-B104. (b), (c) and (d) g_s_ responding to 0.5 μM ABA treatments in PIP2;5 deregulation lines and their WT plants.

Fig.S3 Stomatal aperture after ABA treatments. (a) and (b) PIP2;5 OE-4. (c) and (d) PIP2;5 OE-13. (e) and (f) *pip2;5* KO. The PIP2;5 OE, *pip2;5* KO and the corresponding WT are indicated by OE, KO and WT, respectively. “−” and “+” indicated with and without ABA treatments, respectively. Treatment time is indicated.

Fig.S4 ROS accumulation in guard cells after ABA treatments. (a) and (b) PIP2;5 OE-4. (c) and (d) PIP2;5 OE-13. (e) and (f) *pip2;5* KO. The PIP2;5 OE, *pip2;5* KO and the corresponding WT plants are indicated by OE, KO and WT, respectively. “−” and “+” indicate with and without ABA treatments, respectively. Treatment time is indicated.

