## Supplemental figures for "Aquaporin mediating stomatal closure is associated with water conservation under mild water deficit"

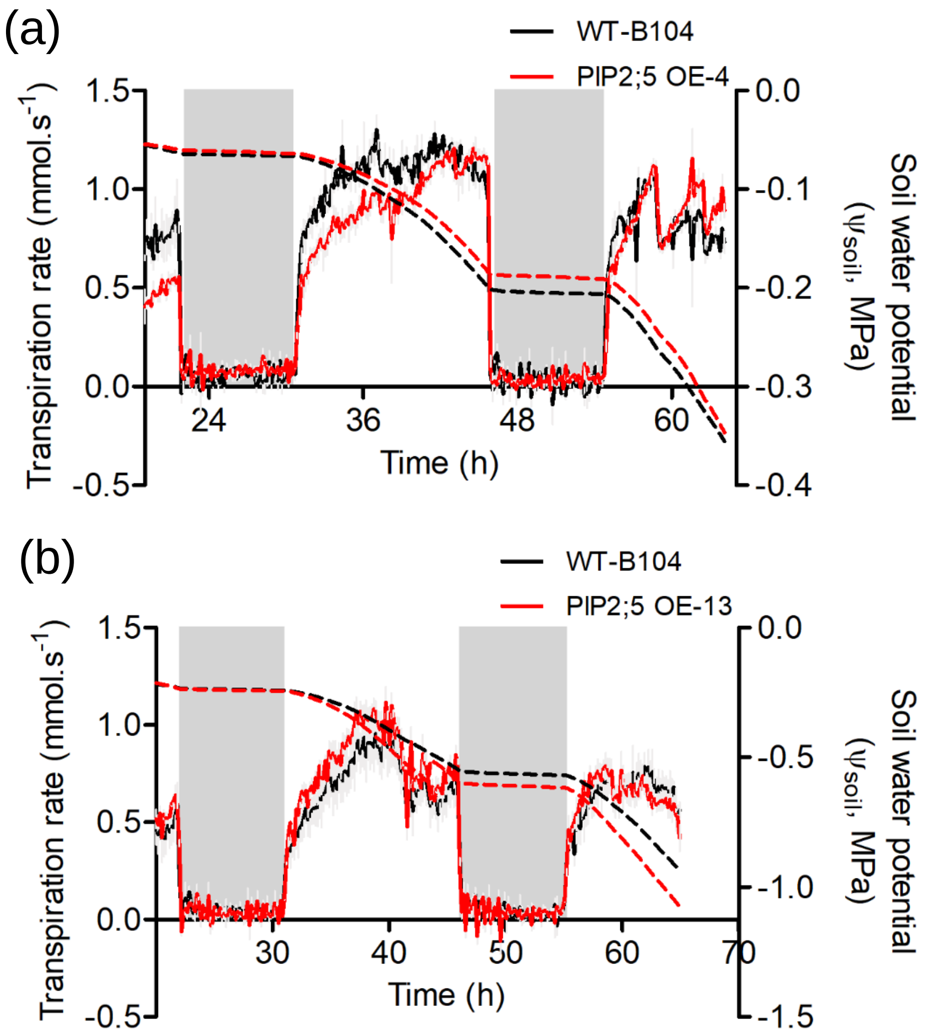


Fig. S1 Transpiration in PIP2;5 OE plants. (a) The transpiration of PIP2;5 OE-4 plants under mild water deficit and severe water deficit conditions. (b) The transpiration of PIP2;5 OE-13 plants under severe water deficit conditions. The dash lines indicate the soil water potential.

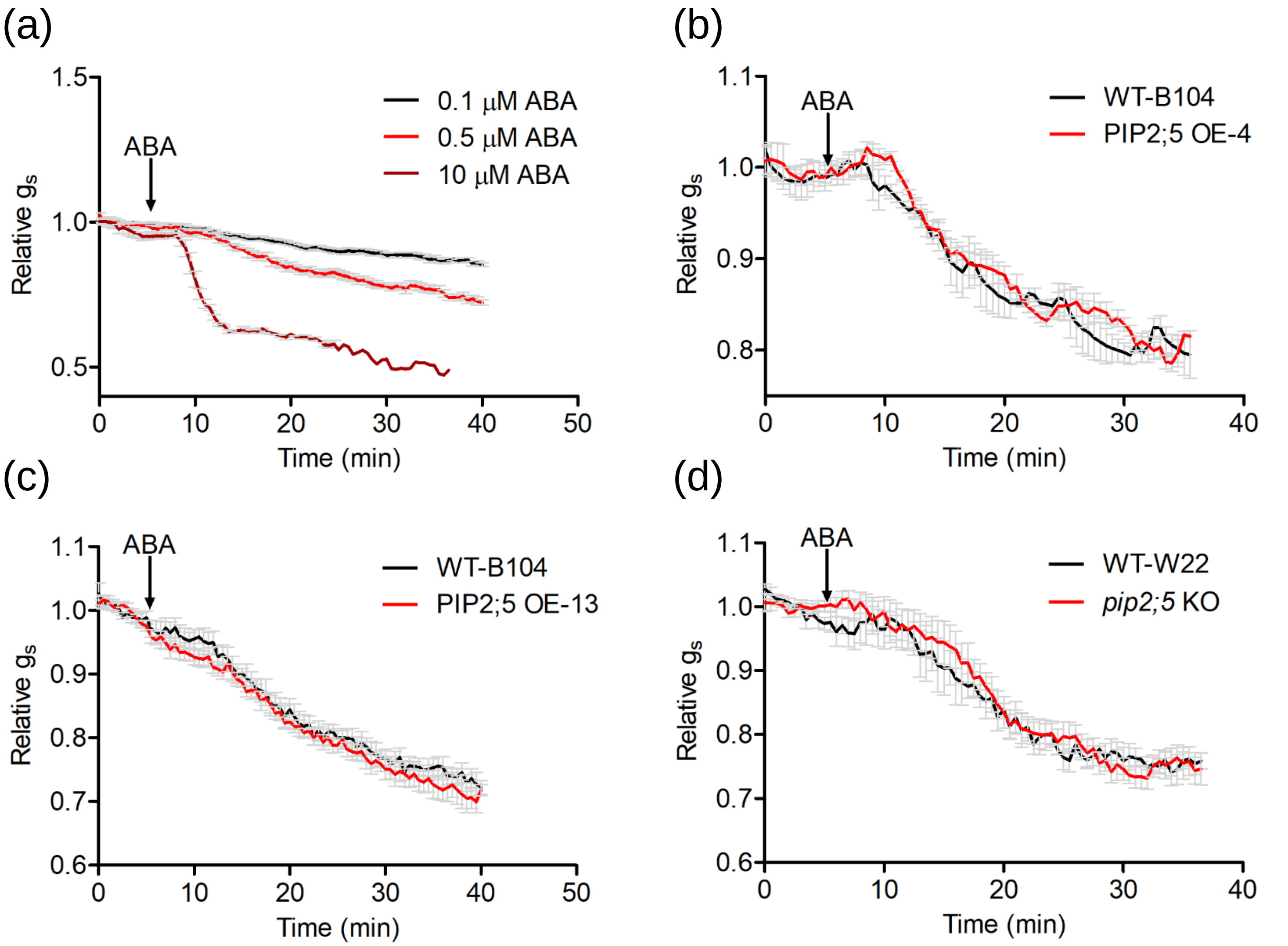


Fig. S2 $g_s$ dynamic after ABA treatment. (a) $g_s$ responding to different ABA concentrations in WT-B104. (b), (c) and (d) $g_s$ responding to 0.5 μM ABA treatments in PIP2;5 deregulation lines and their WT plants.

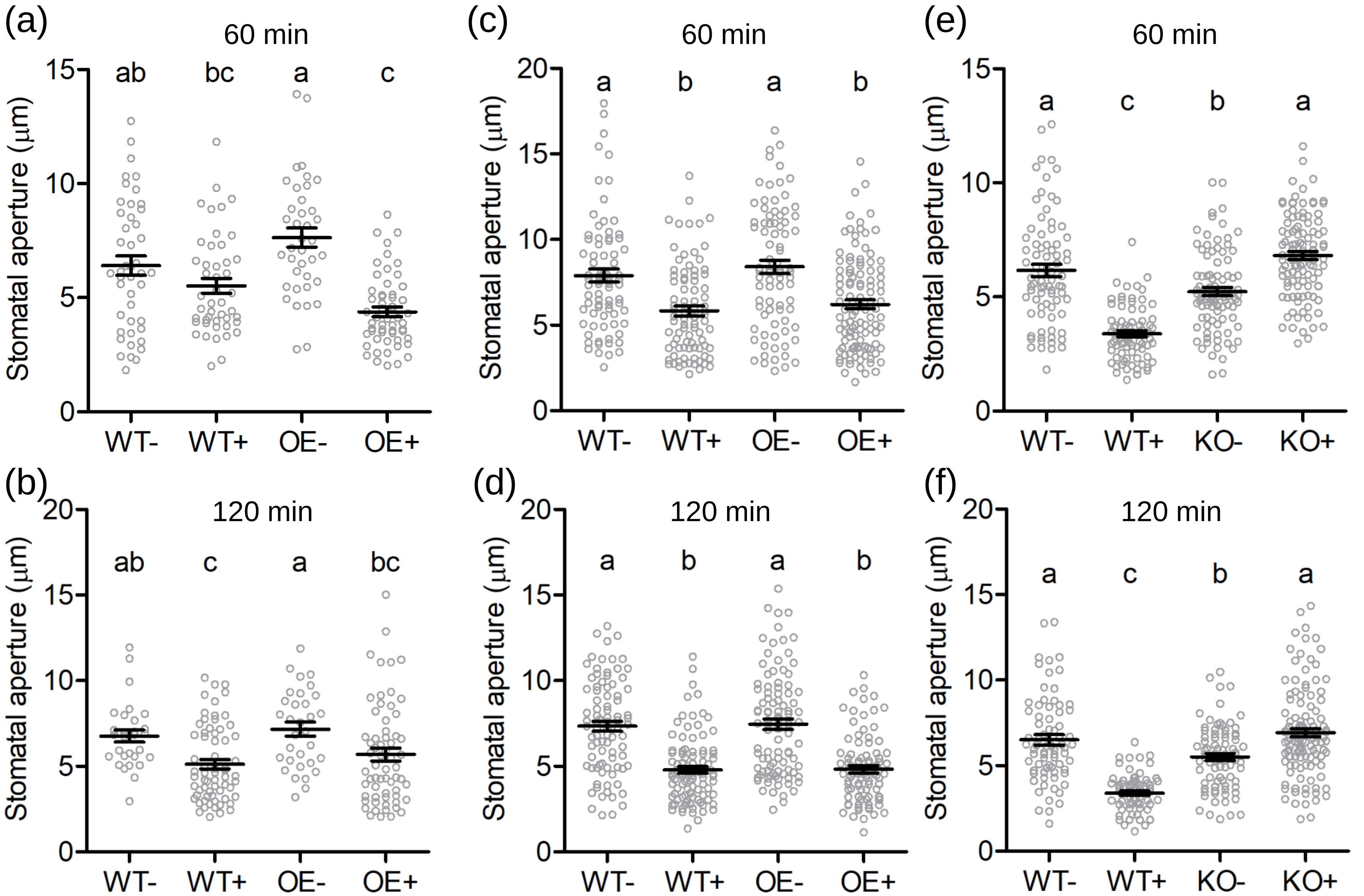


Fig. S3 Stomatal aperture after ABA treatments. (a) and (b) PIP2;5 OE-4. (c) and (d) PIP2;5 OE-13. (e) and (f) *pip2;5* KO. The PIP2;5 OE, *pip2;5* KO and the corresponding WT are indicated by OE, KO and WT, respectively. "-" and "+" indicated with and without ABA treatments, respectively. Treatment time is indicated.

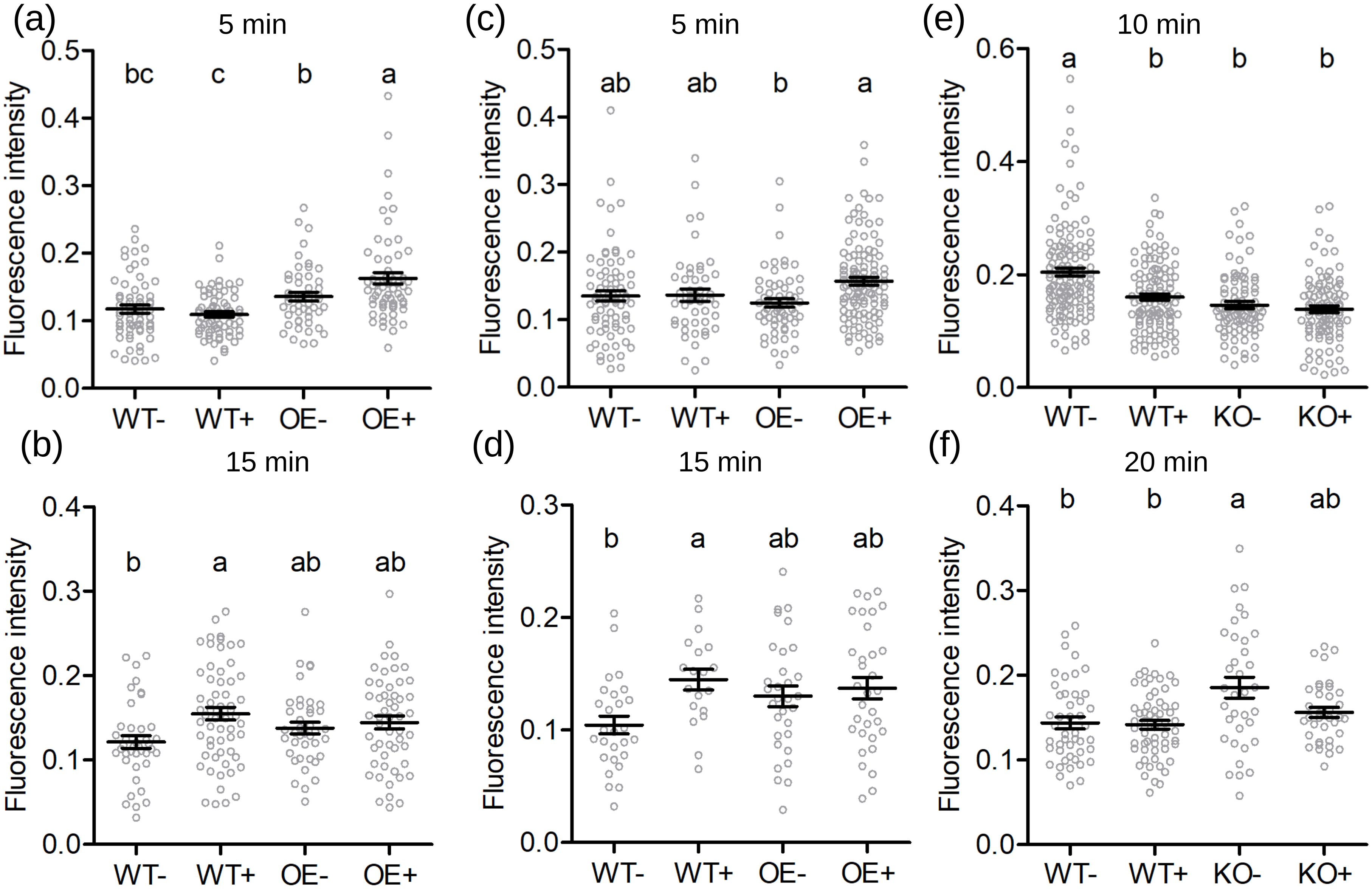


Fig. S4 ROS accumulation in guard cells after ABA treatments. (a) and (b) PIP2;5 OE-4. (c) and (d) PIP2;5 OE-13. (e) and (f) *pip2;5* KO. The PIP2;5 OE, *pip2;5* KO and the corresponding WT plants are indicated by OE, KO and WT, respectively. “-” and “+” indicate with and without ABA treatments, respectively. Treatment time is indicated.
